# AnnapuRNA: a scoring function for predicting RNA-small molecule interactions

**DOI:** 10.1101/2020.09.08.287136

**Authors:** Filip Stefaniak, Janusz M. Bujnicki

**Affiliations:** Laboratory of Bioinformatics and Protein Engineering, International Institute of Molecular and Cell Biology, Warsaw, Poland; Institute of Molecular Biology and Biotechnology, Faculty of Biology, Adam Mickiewicz University, Poznan, Poland

## Abstract

RNA is considered as an attractive target for new small molecule drugs. Designing active compounds can be facilitated by computational modeling. Most of the available tools developed for these prediction purposes, such as molecular docking or scoring functions, are parametrized for protein targets. The performance of these methods, when applied to RNA-ligand systems, is insufficient. To overcome these problems, we developed AnnapuRNA, a new knowledge-based scoring function designed to evaluate RNA-ligand complex structures, generated by any computational docking method. We also evaluated three main factors that may influence the structure prediction, i.e., starting conformer of a ligand, the docking program, and the scoring function used. We applied the AnnapuRNA method for a *post-hoc* study of the recently published structures of the FMN riboswitch. Software is available at https://github.com/filipspl/AnnapuRNA

**Author Summary:** Drug development is a lengthy and complicated process, which requires costly experiments on a very large number of chemical compounds. The identification of chemical molecules with desired properties can be facilitated by computational methods. A number of methods were developed for computer-aided design of drugs that target protein molecules. However, recently the ribonucleic acid (RNA) emerged as an attractive target for the development of new drugs. Unfortunately, the portfolio of the computer methods that can be applied to study RNA and its interactions with small chemical molecules is very limited. This situation motivated us to develop a new computational method, with which to predict RNA-small molecule interactions. To this end, we collected the information on the statistics of interactions in experimentally determined structures of complexes formed by RNA with small molecules. We then used the statistical data to train machine learning methods aiming to distinguish between RNA-ligand interactions observed experimentally and other interactions that can be observed in theoretical analyses, but are not observed in nature. The resulting method called AnnapuRNA is superior to other similar tools and can be used to predict preferred ligands of RNA molecules and how RNA and small molecules interact with each other.

## Introduction

Ribonucleic acids (RNAs) play a pivotal role in many cellular processes such as transmission of genetic information, sensing and communicating responses to cellular signals, and even catalysis of chemical reactions [1]. These processes are often modulated by other molecules, such as ions or small molecules, making RNA an attractive therapeutic target for new drugs [2] [3] [4]. A well-studied and clinically validated example is the bacterial ribosome, a molecular target for many antibiotics [5]. Riboswitches, a class of regulatory RNA structural elements embedded in untranslated regions of mRNAs, comprise another interesting group of RNA targets. These structural elements can directly bind a ligand to regulate gene function without the need for protein cofactors [6][7]. Riboswitches are common in bacterial cells and rarely occur in eukaryotic cells, which makes them suitable targets for new antibacterial drugs. One well-studied example is the FMN riboswitch, which controls expression of genes required for biosynthesis and transport of riboflavin (vitamin B2) in bacteria [8]. Several small-molecule inhibitors of the FMN riboswitch with proven antibacterial properties have been identified. Some of these small molecule inhibitors include compounds such as roseoflavin [9,10] or 5FDQD [11], which are structurally similar to the natural ligand, as well as structurally dissimilar ligands having a different chemotype, for example, ribocil and its derivatives [12]. Apart from riboswitches, other medically-relevant RNAs include viral RNAs (dimerization initiation site of HIV-1 RNA [13]), self-splicing group I introns (e.g., an inhibitor for the *td* intron RNA [14]), group II introns (e.g., inhibitors of group IIB intron splicing [15]), and viral ribozymes (e.g, hepatitis D virus ribozyme inhibition by aminoglycosides [16]) (for details, see [17]).

Elucidating the role of RNA-ligand interactions and design of new RNA-binding molecules can be facilitated by analyzing three dimensional (3D) structures of the RNA of interest or its complex with a ligand. Unfortunately, experimental determination of 3D structures is an intensive task and often has to be supported by computational modeling [18] or performed entirely *in silico* (for review on methods of modeling of ribonucleic acid–ligand interactions see: [19]). These limitations,, together with an increasing interest in RNA as a target for therapeutic intervention, highlight the need to develop new methods for predicting the 3D structure of RNA-ligand complexes.

One of the most widely used computational methods used to predict the 3D structures of macromolecules with ligands is molecular docking [20,21]. Many of the currently-available docking programs were initially designed for protein-protein or protein-ligand docking (e.g., as AutoDock [22], AutoDock Vina [23], ICM [24], or iDock [25]) while some of them were later adapted or reparameterized to enable RNA-ligand docking, where RNA is specified as the receptor (Dock6, [26], ICM [27], or AutoDock [28]). A few programs were designed and optimized specifically for docking ligands to RNA. MORDOR allows for both ligand and receptor flexibility and uses a scoring function that estimates the total energy of the complexes and consists of several terms (electrostatic, van der Waals, dihedral angle, torsion angle, bond, and Urey−Bradley) [29]. rDock (formerly: RiboDock) consists of a genetic algorithm stochastic search engine as a pose generator, and intermolecular scoring function validated against protein and RNA targets [30,31]. The main component of this scoring function is a van der Waals potential, an empirical term for attractive and repulsive polar interactions, and an optional desolvation potential that combines a weighted solvent accessible surface area approach. rDock program can also be used for rescoring poses generated by an external tool.

In a portfolio of methods facilitating prediction of 3D RNA-ligand structures, there are also standalone scoring functions specific for RNA-ligand complexes, intended to be used for rescoring of models generated by molecular docking. A separate group of such methods consists of statistical potentials (statistical scoring functions, or knowledge-based potentials), which are derived from an analysis of experimentally solved structures. Pfeffer and Gohlke developed a scoring function named DrugScore^RNA^, which employs a distance-dependent potential calculated on the basis of contacts between ligand and receptor atoms [32]. Both interacting partners are in all-atom representation using Tripos atom types. Similarly, a KScore scoring function, described by Zhao *et al.*, is also a distant-dependent potential, which in addition to the standard set of Tripos atom types, uses an extended atom type set to characterize the metal-ligand and water-ligand interactions better. It likewise defines a special atom-typing scheme for nucleic acids, and it was parameterized on ligand complexes with proteins (2422 complexes), DNA (300 complexes), and RNA (97 complexes). Yan and Wang described a distant-dependent scoring function termed SPA-LN, where Tripos atom types are used for RNA and ligand representation [33]. During the optimization of parameters, they focused not only on recapitulating the near-native pose of the ligand but also on the affinity estimation. Improving the affinity predictions was a goal of Chen *et al*., authors of iMDLScore1 and iMDLScore2 functions [34]. They optimized AutoDock4.1 scoring terms using multilinear regression methods and binding affinity data for 45 RNA complexes. Our group developed LigandRNA - an anisotropic distance- and angle-dependent statistical potential [35]. It was derived from a diversified set of 251 experimentally solved RNA-ligand complexes and used all-atom representation with Tripos atom types. It was shown to be superior to described earlier scoring functions in terms of accuracy of finding a near-native conformation of ligands. Chhabra *et al.* in the RNAPosers used a distant dependent fingerprint to describe a binding pose of a ligand in RNA binding pocket [36]. Data derived from 80 experimentally solved RNA-ligand complexes were used to train a machine-learning algorithm, which was then used to rank docking poses. Similarly to the programs described above, RNAPosers uses an all-atom representation with Tripos atom types.

In this study, we describe AnnapuRNA, a new knowledge-based scoring function designed to evaluate RNA-ligand complex structures, generated by any computational docking method. We also present a benchmark in which we compare AnnapuRNA with other available scoring functions. We present a case study in which we used molecular docking in combination with the AnnapuRNA scoring function to predict a recently published structure of an FMN riboswitch co-crystallized with a new small-molecule inhibitor. Our method successfully predicted the structure of this complex based on the previously published structure of FMN in complex with a different ligand, as well as the low-resolution apo structure of this RNA.

AnnapuRNA is freely available to the scientific community at https://github.com/filipspl/AnnapuRNA.

## Results and discussion

AnnapuRNA is a machine-learning statistical scoring function developed by us for predicting the structure of RNA-ligand complexes with high accuracy. Our program internally uses a coarse-grained representation for both interacting partners - RNA and small molecule ligands. Using coarse-grained representation has been proven successful for simulating various biomolecular systems [37], including folding of the RNA molecule by the SimRNA program, also developed by our group. We derived machine learning models expressing the probability of RNA-ligand interactions, based on the interaction data collected from experimentally solved complexes extracted from the PDB database. AnnapuRNA can be used to identify native-like poses of ligands from the pool of poses generated through molecular docking experiments. It supports outputs from multiple docking programs (such as rDock, iDock, AutoDock Vina, Dock6, among others).

To evaluate the accuracy of AnnapuRNA in finding near-native structures of RNA-ligand complexes, we carried out extensive benchmarking. We tested three main factors that are likely to influence the outcome of the structure prediction procedure: the starting conformer of a ligand, the docking program, and the scoring function used (see Fig 1). Finally, we used molecular docking for conducting *post-hoc* studies of the recently published structures of the FMN riboswitch.

**Fig 1.**
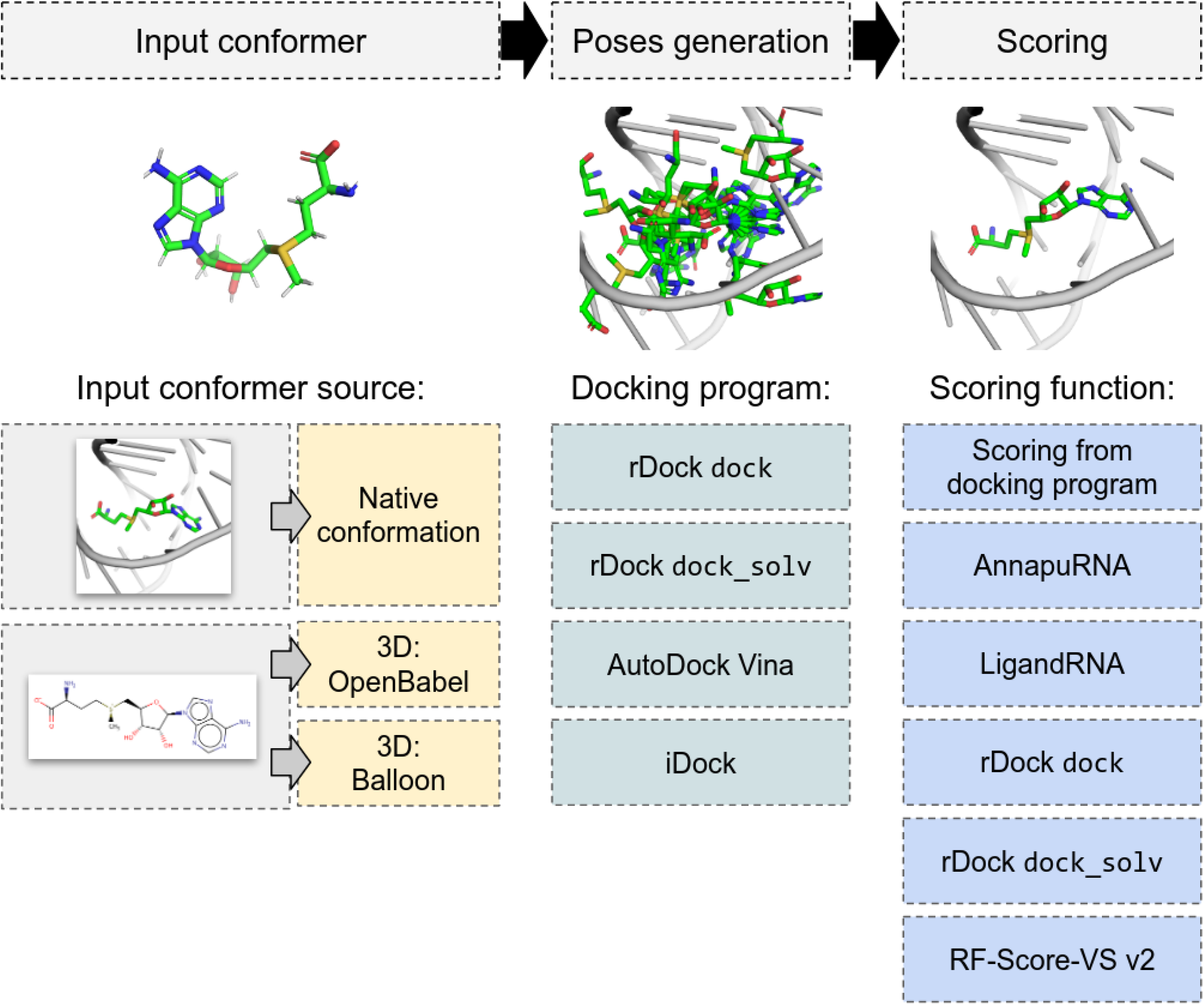
Typical stages of molecular docking applied for interaction predictions for RNA-ligand complexes and methods tested within this publication. The first stage is ligand conformer generation (either using the native conformation extracted from experimentally solved structure or generation of 3D structure), which is followed by molecular docking, and scoring of generated poses.

### Comparison of AnnapuRNA with other scoring methods

To compare the performance of AnnapuRNA scoring functions with methods developed earlier, we used a test set described earlier by Philips *et al*. [35]. AnnapuRNA models were trained on the full “2013” and “2016” training sets. In this analysis, we used a pose of a ligand extracted from the experimentally solved structure as an input conformation and the rDock program with dock desolvation potential for docking. For each complex, we generated a set of 100 poses and applied an external scoring function for rescoring. We tested nine rescoring methods: RF-Score-VS v2 (Random Forest-based scoring function for Virtual Screening, trained for proteins, [38]), rDock scoring functions with two desolvation potentials: dock and dock_solv (these potentials, implemented in rDock docking program, consist of a van der Waals potential, an empirical term for attractive and repulsive polar interactions, and an optional desolvation potential), LigandRNA (a knowledge-based potential derived from ligand-binding sites in the experimentally solved RNA– ligand complexes, obtained using the inverse Boltzmann scheme; this scoring function was used in two versions: the one described in the original publication, called “2013”, and the second, with an updated potential, called “updated” [35], and AnnapuRNA described herein (variants based on two machine learning methods: DL and *k*NN, with two sets of training data: “2013” and “2016”). Regrettably, we were unable to run the DrugScore^RNA^ scoring function, despite contacting the authors and obtaining the source code [32]. Also, the authors of the recently published SPA-LN scoring function [33] were unable to provide us with the working copy of their program nor give access to the dedicated web server. Results expressed as RMSD of the best-selected pose to the reference pose (*S*(1)), *S*(3), and *S*(5) are summarized in the Fig 2, Supplementary Fig S1, and Supplementary Table S8 in S1 Text. In addition to the results of rescoring, we included data for the median RMSD values obtained during docking (this serves as a negative control) and minimum RMSD values (a positive control).

**Fig 2.**
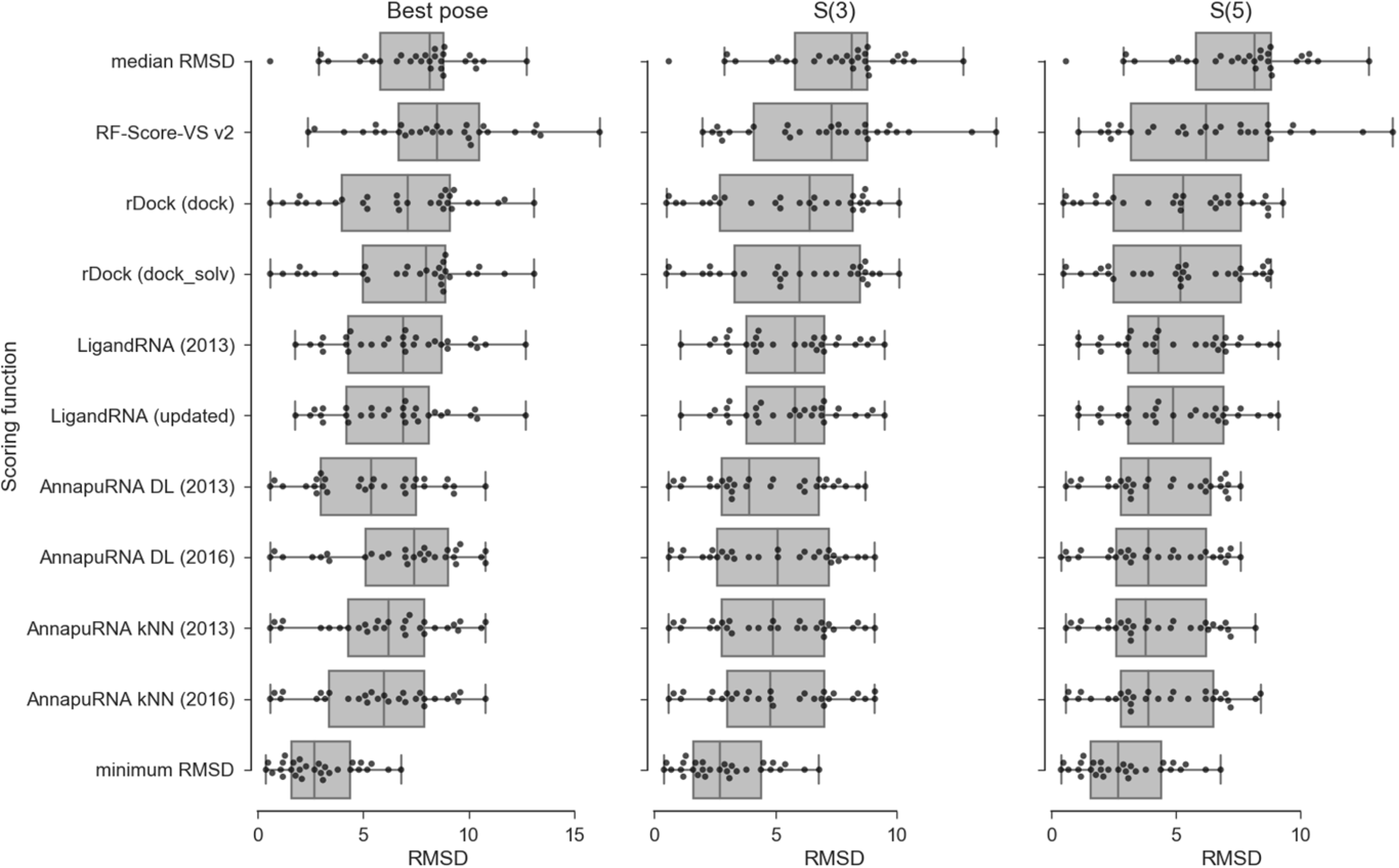
Comparison of the performance of nine scoring functions, expressed as the RMSD between the best-scored pose and the reference pose (left panel), best among the top three scored poses (*S* (3), middle panel), and best among the top five scored poses (*S* (5), right panel). Additional rows represent the median and minimal values of RMSD obtained during docking. Each dot represents one complex from the testing set. Docking was performed using rDock with dock desolvation potential with the native conformation of a ligand as an input.

The average values for all three metrics: *S*(1), *S*(3), and *S*(5) indicate that the AnnapuRNA scoring functions are, in most cases, superior (i.e., having lower values of *S*(n)) to the methods developed previously. For the most strict criteria, which is the average RMSD of the best-scored poses (*S*(1)), the best performing methods were AnnapuRNA version DL trained on the “2013” dataset and AnnapuRNA version *k*NN trained on “2016” dataset (*S*(1) equal to 5.39 and 5.76, respectively). Among the LigandRNA variants, the updated version of the potential (*S*(1) = 6.37) performed better than the older version. Among the rDock variants, the dock desolvation potential (*S*(1) = 6.81) was found to be better.

In this comparison, AnnapuRNA generally outperforms all the other methods, with the only exception of AnnapuRNA DL trained on the “2016” dataset that is outperformed by our previously-built method, LigandRNA in selecting the single best-RMSD pose. The overall best-performing potential is AnnapuRNA DL-2013.

The performance of the ‘protein-focused’ scoring function, RF-Score-VS v2, was found to be worse than all methods tested. Both median and average values for all the three metrics tested are high for this method, with a very high variation in the results. For example, *S*(3) for this method ranges from 2.0 to 14.1, with a standard deviation of 3.2. These findings show that the development of an RNA-specific scoring function is necessary for advancing RNA-ligand docking and modeling of RNA-ligand interactions.

The comparison of LigandRNA and AnnapuRNA, scoring functions developed in our group (both variants trained on structures available in 2013, i.e., the “2013” versions), indicates that for most metrics used AnnapuRNA (the new method) is superior to LigandRNA. This means that based on the same subset of experimentally solved structures available for method training, we obtained more accurate models of interactions using the approach described in this article. Moreover, AnnapuRNA is 5-6 times faster than LigandRNA, which is an important factor taken into consideration when processing large datasets (see: Table S13 and Fig S5 in the S1 Text). The main differences in the design of these programs include the coarse-grained representation of molecules and the use of a rich set of interaction descriptors combined with machine learning methods in AnnapuRNA, versus full atom representation, two features describing interactions, and the inverse Boltzmann scheme for deriving potential in LigandRNA.

### Docking programs with selected scoring functions

An important factor to consider in docking is the ability to generate the RNA-ligand pose that resembles the reference structure. To identify the best approach for pose generation, we compared the performance of four molecular docking methods in combination with several scoring functions. For pose generation, we tested rDock (with two desolvation potentials: dock and dock_solv), AutoDock Vina, and iDock. Only rDock is specifically designed for docking ligands to RNA or protein targets, while iDock and Autodock Vina are parameterized only for protein targets. The goal of this analysis was to find the docking program that generates a set of poses with at least one pose having low RMSD to the reference structure, using the ligand conformer taken from the reference structure.

The best pose generator was found to be rDock with dock_solv desolvation potential. The average minimum RMSD for this method was 2.3 Å, while for the dock_solv potential it was slightly higher - 2.6 Å (see: Fig 3, and Table S9 in S1 Text for details). The RMSD values for Autodock Vina and iDock were remarkably higher - 7.1 Å and 6.5 Å, respectively. For two other RNA-ligand complexes, Autodock Vina and iDock generated solutions with remarkably poor RMSD (19.1 Å and 32.3 Å, respectively). This observation is not surprising - in general, programs designed specifically for RNA-ligand docking generate better RNA-ligand complex models, while the two docking programs, trained for protein-ligand docking, generate conformations more distant to the reference structure. On the other hand, rDock was unable to generate any solutions for four complexes in our dataset, for which AutoDock Vina and iDock successfully generated poses, albeit with high RMSD to the native structure. Also, in two cases, AutoDock Vina and iDock were able to generate poses with an RMSD equal to or better than those generated by rDock. These findings imply that in some ‘difficult’ cases, protein-based docking programs may be used to complement the RNA-based methods to generate poses in RNA-ligand docking.

**Fig 3.**
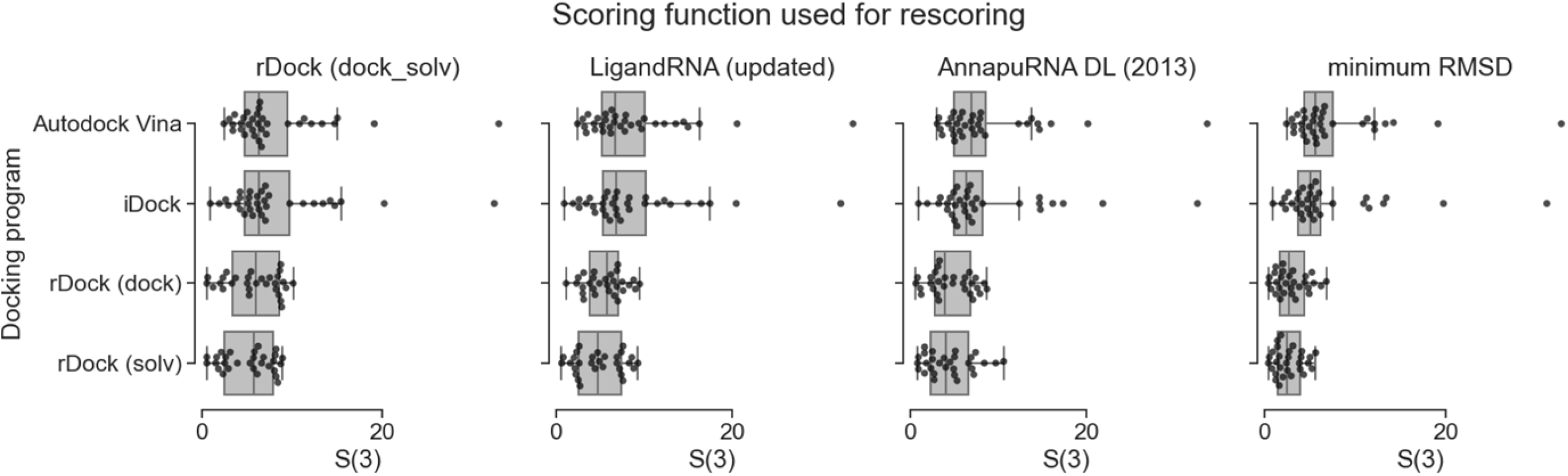
Comparison of the performance of three scoring functions (rDock dock_solv, LigandRNA, and AnnapuRNA) expressed as RMSD to the reference pose of best in top three scored poses (*S* (3)), calculated for four docking programs. The fourth column represents the best (lowest) RMSD obtained during docking for each program. Each dot represents one complex from the testing set. Docking was performed with the native conformation of a ligand as an input.

*S*(3) values for all scoring functions tested are the best for poses generated by rDock. Among all combinations of pose-generating programs with scoring functions, the best performance was achieved for rDock dock_solv in combination with AnnapuRNA scoring functions (see: Table S10 in S1 Text for RMSD values for all complexes in a testing set and Fig S2 in S1 Text for *S*(1) and *S*(5) values distribution).

### Conformer generation methods

In the assessment of RNA-ligand pose generators, we use the pose extracted from the experimentally solved structure. However, in real-life cases, the ‘true’ conformation of the ligand is unknown, and it must be predicted for executing the step of pose generation. Hence, we extended our tests to include a comparison of results obtained from docking ligands whose conformers were generated independently of the known bound conformation, using two programs, OpenBabel and Balloon.

We found that incorporating the native ligand conformation does not provide a significant advantage in RNA-ligand docking. The differences between the results obtained with the experimentally observed and predicted poses were minimal. The best sets of poses (i.e., the pools of poses containing a pose with the lowest RMSD to the reference structure) were generated when the Balloon program was used as a conformer generator (the average minimal RMSD 2.86 Å), minimally better than the results obtained for the native poses (2.90 Å, see Fig 4). This means that the tested docking programs can sample a conformational space of ligands quite effectively and find the desired RNA-ligand interactions regardless of the starting ligand conformation. This effect is observed regardless of the scoring function used to identify the final solutions (see also Fig S3 in S1 Text for *S*(1) and *S*(5) values distribution). From this analysis, we conclude that the key elements of the docking pipeline are the choice of the docking program and the scoring function, while the input conformation of a ligand plays a less important role.

**Fig 4.**
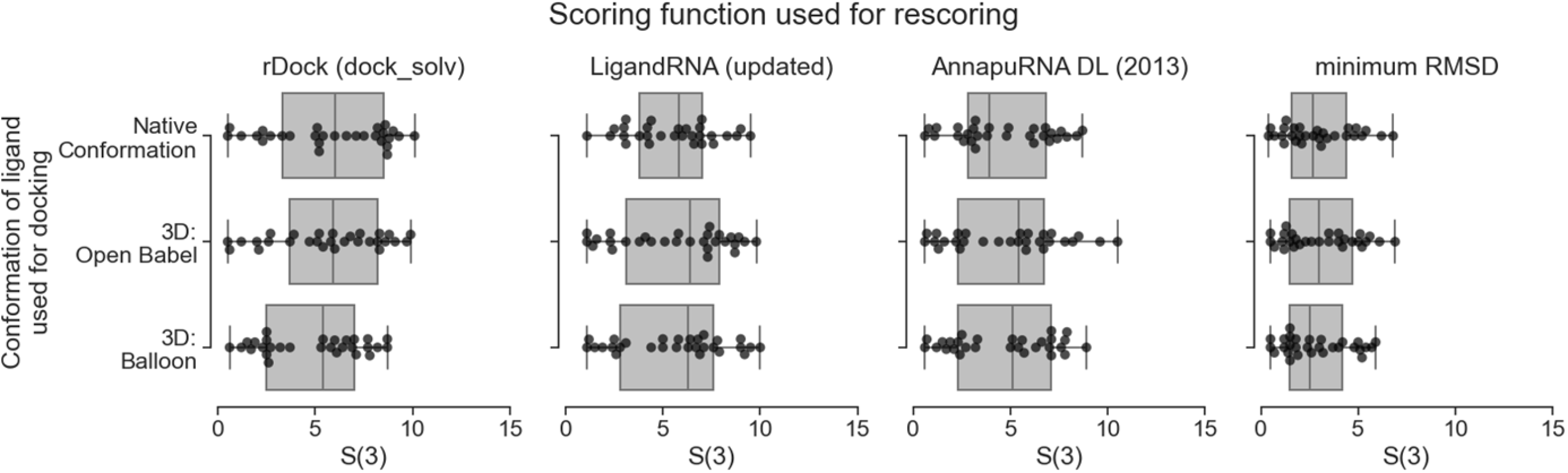
Comparison of the performance of three scoring functions, expressed as a RMSD of the best-scored pose to the reference pose of best in top three scored poses (*S*(3)), calculated for three conformer generation methods. The fourth column represents the best (lowest) RMSD obtained during docking for each program. Each dot represents one complex from the testing set. Docking was performed using rDock with dock desolvation potential.

Analysis of the statistical significance of our results reveals that although the average and median performance *S*(n) of AnnapuRNA is better than that of other programs, these differences are not always statistically significant. For example, for the median value of the best RMSD for rDock docking program with dock_solv desolvation potential, the AnnapuRNA *k*NN (2013) method is significantly different (better) than all other scoring functions (*p* values ranges from <0.001 for the RF-Score-VS v2, to 0.033 for the updated version of LigandRNA) except the LigandRNA (2013, *p* = 0.053; see Table 1). Tests performed for the same methods, but for the AutoDock docking program show that the AnnapuRNA method is significantly different from rDock (dock_solv) scoring function (*p* = 0.038), while for others functions the difference is not significant (*p* > 0.05).

**Table 1.** *P*-values obtained in a Wilcoxon signed-rank test comparing the distribution of *S*(1), *S*(3), and *S*(5) values of AnnapuRNA and other docking programs. Docking was performed using native conformation of a ligand as an input. Two-tailed *p*-values ≤ 0.05 are in bold. Data referenced in the main text are underlined.

| | | Wilcoxon signed-rank test $p$-value (two tailed) | | | | | | | | | | | |
| --- | --- | --- | --- | --- | --- | --- | --- | --- | --- | --- | --- | --- | --- |
|  |  | docking program:<br>rdock (dock_solv) |  |  | docking program:<br>rdock (dock) |  |  | docking program:<br>AutoDock Vina |  |  | docking program:<br>iDock |  |  |
| AnnapuRNA<br>variant | Compared<br>method | $S(1)$ | $S(3)$ | $S(5)$ | $S(1)$ | $S(3)$ | $S(5)$ | $S(1)$ | $S(3)$ | $S(5)$ | $S(1)$ | $S(3)$ | $S(5)$ |
| AnnapuRNA<br>DL (2013) | LigandRNA<br>(2013) | 0.201 | 0.601 | 0.296 | 0.104 | 0.095 | 0.198 | 0.933 | 0.939 | 0.444 | 0.719 | 0.964 | 0.863 |
|  | LigandRNA<br>(updated) | 0.162 | 0.455 | 0.286 | 0.211 | 0.088 | 0.122 | 1.000 | 0.977 | 0.390 | 0.478 | 0.964 | 0.713 |
|  | rDock (dock) | **0.009** | **0.042** | **0.036** | **0.032** | **0.024** | **0.044** | 0.381 | **0.019** | 0.206 | **0.009** | **<**<br>**0.001** | **0.001** |
|  | rDock<br>(dock_solv) | **0.005** | 0.057 | 0.056 | **0.016** | **0.014** | **0.027** | 0.171 | **0.035** | 0.087 | 0.125 | 0.128 | **0.027** |
|  | RF-Score-VS<br>v2 | **0.001** | **0.001** | **0.002** | **<**<br>**0.001** | **0.001** | **0.002** | 0.173 | 0.117 | 0.206 | 0.365 | 0.094 | **0.016** |
| AnnapuRNA<br>DL (2016) | LigandRNA<br>(2013) | 0.465 | 0.811 | 0.078 | 0.819 | 0.429 | 0.140 | 0.540 | 0.484 | 0.848 | 0.339 | 0.755 | 0.636 |
|  | LigandRNA<br>(updated) | 0.313 | 0.984 | 0.073 | 0.475 | 0.394 | 0.085 | 0.597 | 0.513 | 0.768 | 0.194 | 0.755 | 0.567 |
|  | rDock (dock) | **0.023** | 0.136 | **0.008** | 0.889 | 0.053 | **0.008** | 0.799 | 0.069 | **0.033** | 0.050 | **<**<br>**0.001** | **0.002** |
|  | rDock<br>(dock_solv) | **0.014** | 0.368 | **0.022** | 0.647 | 0.062 | **0.009** | 0.407 | 0.079 | **0.011** | 0.241 | 0.078 | **0.025** |
|  | RF-Score-VS<br>v2 | **0.001** | **0.006** | **0.001** | **0.011** | **0.001** | **0.002** | **0.033** | 0.264 | 0.762 | 0.130 | **0.049** | **0.047** |
| AnnapuRNA<br>kNN (2013) | LigandRNA<br>(2013) | <u>0.053</u> | 0.542 | 0.636 | 0.200 | 0.465 | 0.082 | <u>0.459</u> | 0.867 | 0.809 | 0.468 | 0.442 | 0.316 |
|  | LigandRNA<br>(updated) | **0.033** | 0.469 | 0.619 | 0.456 | 0.404 | **0.048** | <u>0.346</u> | 0.886 | 0.809 | 0.264 | 0.442 | 0.183 |
|  | rDock (dock) | **0.004** | **0.039** | **0.050** | 0.361 | 0.068 | **0.009** | <u>0.107</u> | **0.028** | 0.135 | 0.052 | **<**<br>**0.001** | **<**<br>**0.001** |
|  | rDock<br>(dock_solv) | **0.002** | 0.086 | 0.121 | 0.264 | 0.072 | **0.008** | **0.038** | **0.039** | 0.061 | 0.144 | **0.017** | **0.001** |
|  | RF-Score-VS<br>v2 | **<**<br>**0.001** | **0.001** | **0.005** | **0.003** | **0.001** | **0.001** | <u>0.373</u> | 0.245 | 0.334 | 0.166 | 0.217 | 0.472 |
| AnnapuRNA<br>kNN (2016) | LigandRNA<br>(2013) | 0.130 | 0.538 | 0.363 | 0.118 | 0.330 | 0.217 | 0.952 | 0.808 | 0.438 | 0.145 | 0.666 | 0.690 |
|  | LigandRNA<br>(updated) | 0.104 | 0.433 | 0.349 | 0.266 | 0.297 | 0.131 | 0.980 | 0.808 | 0.386 | 0.068 | 0.666 | 0.556 |
|  | rDock (dock) | **0.011** | 0.079 | **0.044** | 0.127 | 0.182 | **0.024** | 0.260 | **0.021** | 0.327 | 0.108 | **0.001** | **0.001** |
|  | rDock<br>(dock_solv) | **0.007** | 0.136 | 0.079 | 0.097 | 0.072 | **0.028** | 0.170 | **0.028** | 0.199 | 0.260 | 0.050 | **0.024** |
|  | RF-Score-VS<br>v2 | **0.001** | **0.001** | **0.004** | **0.001** | **0.003** | **0.002** | 0.098 | 0.309 | 0.119 | **0.042** | 0.153 | 0.059 |

For a summary of all combinations of ligand preparation methods, docking programs and scoring functions tested, see Table S11 in S1 Text.

In the current version of AnnapuRNA, we implemented three optional steps for the post-processing of poses from docking or rescoring, namely clustering, centroid calculation, and local optimization. For details, see: S1 Text, chapter “post-processing of docking poses”.

### Selected examples from the test set

Fig 5 shows redocking results for three selected cases from the testing set, which compares the AnnapuRNA scoring function (presented in this work and found to outperform other methods) with the runner-up, namely the rDock scoring function. We choose three RNA-ligand complexes with small molecules displaying various degrees of flexibility - rigid, bicyclic THF fragment (Fig 5A), semi-rigid paromomycin, containing four rigid, heterocyclic rings connected by rotatable oxygen linkers (Fig 5B), and a flexible, aliphatic argininamide chain (Fig 5C). We expected that with increased flexibility of the ligand, a more diverse set of conformations (with possibly higher RMSD range) would be generated by the docking program and thus it would be increasingly difficult to determine the near-native pose for the scoring function. In all three cases, the AnnapuRNA scoring function selected the pose closer to the reference one than the rDock scoring function. Also, RMSD vs score plots reveal that energy funnels for AnnapuRNA have a more linear shape, while for rDock the points are more uniformly scattered across the plot. This is an interesting observation, taking into account the fact that RMSD values were not used directly to develop the machine learning models of interactions. This could also mean that the AnnapuRNA score could be correlated with the RMSD and then potentially used for the assessment of RMSD values for a given pose. This hypothesis is supported by the results of analysis of three correlation coefficients (Spearman's rank correlation coefficient, Pearson correlation coefficient, and Kendall rank correlation coefficient) for RMSD-score values. For docking with rDock (in both variants), the mean correlation coefficient is higher (which translates to a better correlation between score and the RMSD) for the AnnapuRNA compared to other scoring functions. For example, the mean Spearman's rank correlation coefficient for rDock (dock), LigandRNA (2013) and AnnapuRNA (*k*NN, 2013) is 0.117, 0.244, and 0.333, respectively. For the updated version of our potential, these values are even higher, reaching values of 0.345 and 0.336 for DL and *k*NN, respectively. This means that the score calculated by AnnapuRNA correlates better with RMSD than the score of other scoring functions tested (for details, see Table S14 and S6 in S1 Text).

**Fig 5.**
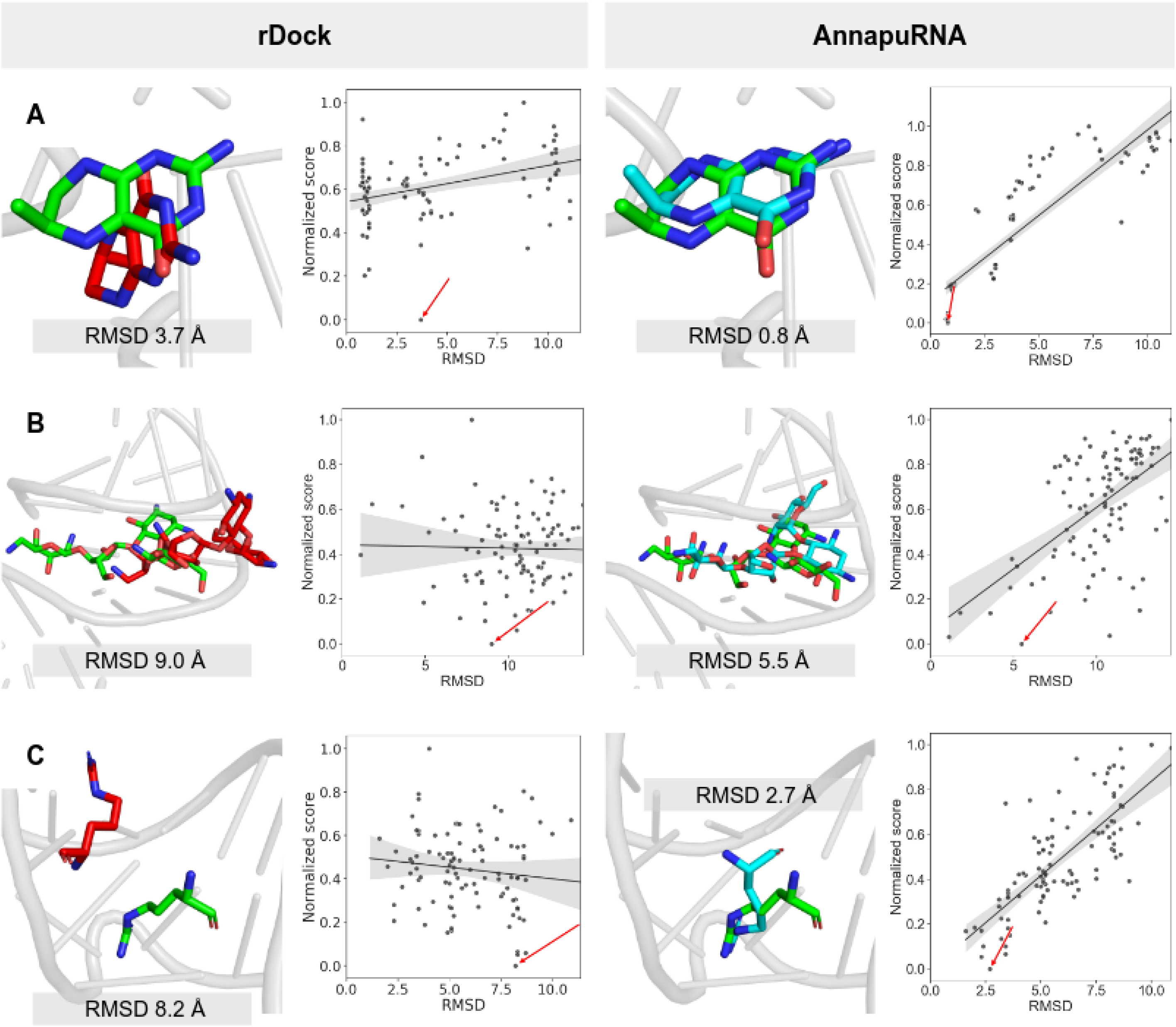
Selected docking solutions for structures from the testing set. Best pose found by rDock (with dock desolvation potential, left) and AnnapuRNA (DL 2013, right), together with scatter plots of RMSD and normalized score. Scatter plots show a regression line with a 95% confidence, and the top scoring pose according to each method is indicated by a red arrow. (**A**) 3SUX (Crystal structure of THF riboswitch, bound with THF fragment), (**B**) 1FYP (Decoding region A-site in complex with Paromomycin) and (**C**) 1AJU (HIV-2 TAR-argininamide complex). RNA molecules are presented as a gray cartoon, ligands as sticks; reference structure - green, solution found by rDock - red, solution found by AnnapuRNA - cyan. Heteroatoms are colored: O—red, N—blue.

### Case study

One of the most important applications of molecular docking and scoring functions is to predict the binding mode of a small molecule in a binding pocket for a macromolecule of interest - a protein or nucleic acid. In a typical situation, the experimentally solved structure of the macromolecule is used for modeling. In most cases, this structure is available either as a complex with a structurally different ligand or as an apo structure. In these cases, predicting the structure of a new complex is more challenging than in a redocking experiment or when a macromolecule structure solved with a structurally similar ligand is used as a receptor [39]. We performed a set of simulations mimicking this real-life scenario. As a model system, we used a recently published structure of the FMN riboswitch, co-crystallized with a small molecule inhibitor (6DN2, resolution 2.880 Å [40]). Our task was to predict the binding mode of this ligand using two structures of FMN riboswitch described seven years earlier [41]: the apo form (2YIF, resolution 3.298 Å), and a complex with a structurally different ligand (2YIE, resolution 2.941 Å). None of these structures were used for training or testing the AnnapuRNA function. During this test, we compared two scoring functions - rDock and AnnapuRNA (DL 2013). The first experiment involved the redocking of the ligand to the native structure (6DN2). Both scoring functions were able to find a pose that is close to the reference; however, AnnapuRNA performed better than rDock (1.1 Å and 1.7 Å, respectively; Fig 6A).

**Fig 6.**
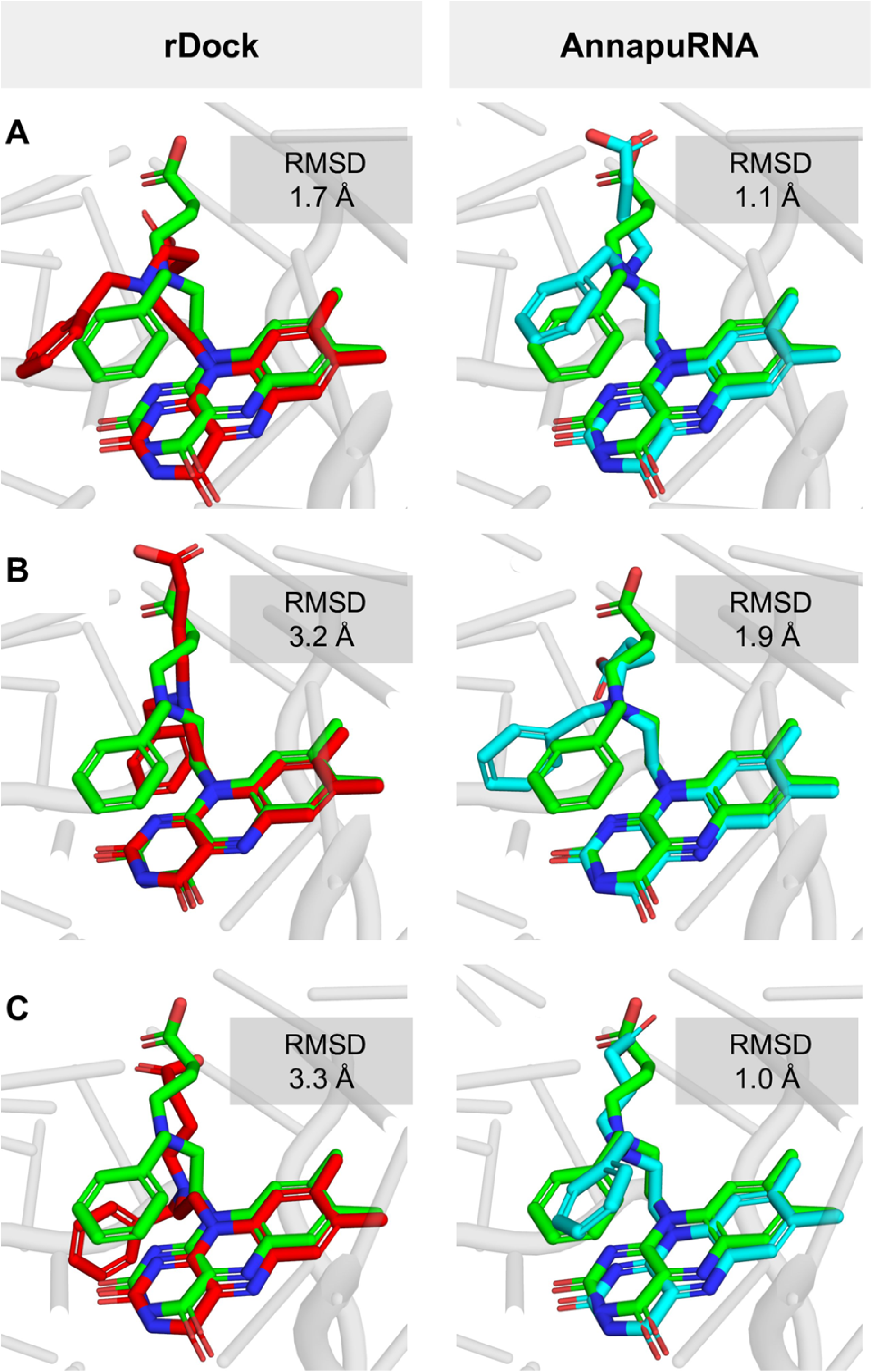
Prediction of the complex of FMN Riboswitch with 4-{benzyl[2-(7,8-dimethyl-2,4-dioxo-3,4-dihydrobenzo[g]pteridin-10(2H)-yl)ethyl]amino}butanoic acid. (**A)**. Redocking experiment - ligand extracted from the crystal structure 6DN2 was redocked to this structure; (**B)**. Ligand extracted from the crystal structure 6DN2 docked to FMN structure solved with a different ligand (2YIE); (**C)**. Ligand extracted from the crystal structure 6DN2 docked to the low-resolution APO FMN structure (2YIF). All RNA structures are superimposed. The reference ligands structures are shown in green. Structures predicted by rDock (dock_solv) are shown in magenta (left), predicted by AnnapuRNA (DL 2013) in light blue (right).

In the second case study, we used the FMN structure solved with a structurally different ligand. Here, AnnapuRNA outperformed the rDock scoring function (1.9 Å and 3.2 Å, respectively; Fig 6B). As could be expected, the most difficult fragment to predict was an aliphatic chain, which, in the case of a pose selected by AnnapuRNA, is closer to the reference structure than that selected by rDock. Also, in the third experiment, where the apo structure of FMN was used for docking, AnnapuRNA selected a pose that is very close to the reference, in contrast to the rDock (1.0 Å and 3.3 Å, respectively; Fig 6C). To our surprise, in the case of AnnapuRNA, docking to the apo structure (Fig 6C) yielded better results than those obtained from docking to RNA structures originally solved with ligands (Fig 6A and Fig 6B). This can be attributed to a high entropic contribution/flexibility of the aliphatic side chain of the ligand and thus the difficulty of the docking program in searching the full conformational space exhaustively during each docking. Nevertheless, for each of these experiments, AnnapuRNA selected a pose having an RMSD of 1.9 Å to the reference, which can be considered as a successful docking (a most commonly used criterion is 2.0 Å to the reference structure [42]). In this experiment, we showed that a combination of molecular docking and rescoring using the AnnapuRNA function may be a valuable tool for predicting the structures of RNA-ligand complexes, even when no good quality RNA structure is available.

## Conclusion

In summary, we provide a novel method, AnnapuRNA, for predicting interactions of RNA with small-molecule ligands. Our tool can be used as a computational workflow together with a docking program to generate a biologically relevant model of the RNA-ligand complex. Tests reported in this manuscript indicate that our new method is superior to other programs of this kind developed to date. AnnapuRNA can be extended to data from molecular dynamics or other molecular simulations. Since AnnapuRNA is a knowledge-based method and depends on the content and quality of databases, its main limitation stems from a relatively low number of experimentally determined RNA-ligand complexes. This number is, however, growing steadily, and we plan to update AnnapuRNA in the future using new RNA-ligand structural data.

## Materials and Methods

Fig 7 summarizes the main steps of the AnnapuRNA method development and use.

**Fig 7.**
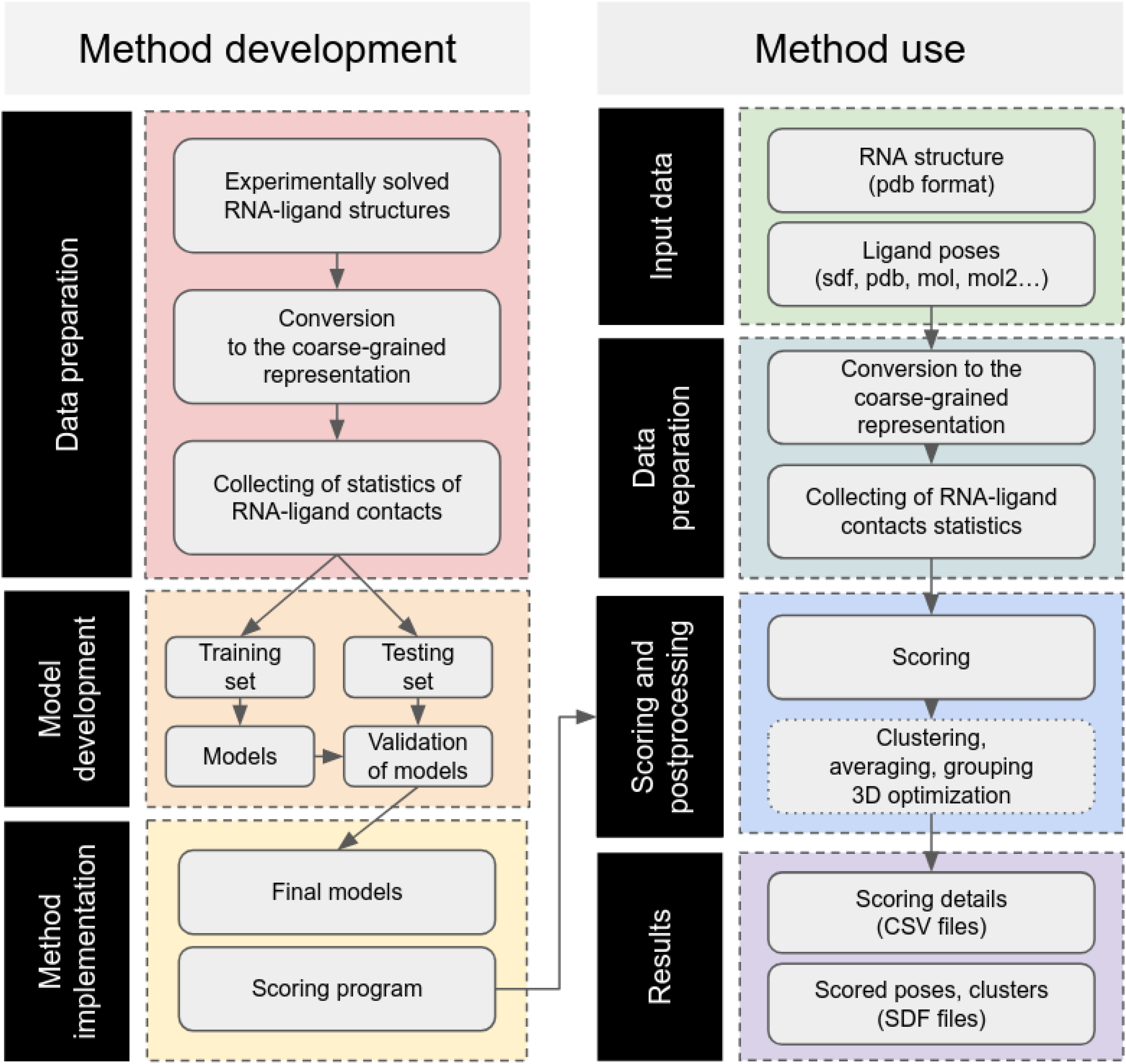
The workflow of AnnapuRNA. The left column represents the method development pipeline, while the right one displays the method usage. Optional post-processing steps are placed in boxes with a dotted border.

### Datasets

Two RNA-ligand structures datasets were prepared. One, called “2013” was based on the structures selected by Philips et al. published until 2013 [35]. The second, called “2016” was updated by additional structures solved and published until 2016.08 (extracted from the PDB database with search criteria: contains RNA: yes; has ligand(s): yes; chemical component type: non-polymer) and selected with the criteria described earlier by Philips at al. (i.e., for RNAs with sequence identity >90%, which are in complex with the same ligand, only the structure with the highest resolution was used; for NMR structures the first model in the file was used; for residues with more than one alternative conformation, the first variant was used; structures containing small molecule ligands closer than 6 Å to any atom other than RNA, water or cation were excluded). Next, for both datasets, RNA structures were preprocessed using the rna-tools software suite (commit b528a29; [43]). Small molecule structures were fetched independently using the RCSB PDB RESTful Web Service interface [44]. Since the AnnapuRNA potential was designed for drug-like compounds, complexes containing ligands with atoms other than C, H, N, O, S, Br, Cl, F, P, Si, B, Se, or lacking C and H atoms were excluded (a molecular formula provided in the xml file fetched from the PDB was used to verify the presence or absence of a given element). To keep only small molecule ligands, we applied the molecular mass criteria with the upper limit of 1000 Da, as defined earlier [45]. This is beyond the most frequently used value of 500 Da [46,47] but allowed us to include larger drug molecules such as oral antibacterial agents, having a substantial number of compounds with the mass in 700-900 Da range [48]. All explicit hydrogens were added to ligand structures with OpenBabel [49].

For each RNA-ligand complex structure, we generated additional near-native poses by redocking the ligand to the RNA receptor. For docking, we used the rDock software, with rigid RNA and flexible ligand, with dock_solv docking protocol and docking radius set to 10. For each complex, 1000 poses were generated and clustered with 2.0 Å RMSD cutoff. We selected a set of poses with RMSD between 0.1 and 1.5 Å to the native ligand pose. The lower value (RMSD < 0.1 Å) was set arbitrarily to exclude poses that are very similar to the native pose. The upper value (RMSD > 1.5 Å) was used earlier to define the threshold of similarity for docking poses [50]. Native and near-native structures were used to derive contact statistics for the “positive” class (native-like solutions). Using the same docking procedure, for each complex, we generated a diversified subset of ligand poses that are different from the native RNA-small molecule structures. For this, we used the RMSD ≥ 4 Å cutoff, which is a doubled value of the commonly used threshold for successful docking [51]. These structures were used to derive contact statistics for the “negative” class (native-unlike solutions).

All ligand structures were converted to a coarse-grained representation with the align-it (version 1.0.3, with noHybrid switch [52]).

### Training and testing set

Both datasets - 2013 and 2016 - were split into two parts (Table 2): the training set, containing 87 and 131 structures respectively, and testing set, with 32 structures common to both datasets (for the list of structures used, see Table S1 in S1 Text). In both cases, the testing set consisted of the structures earlier used for benchmarking RNA-Ligand scoring functions [35]. The optimization of the scoring function was conducted on the training set, and the testing set was not used until the final benchmark.

**Table 2.**
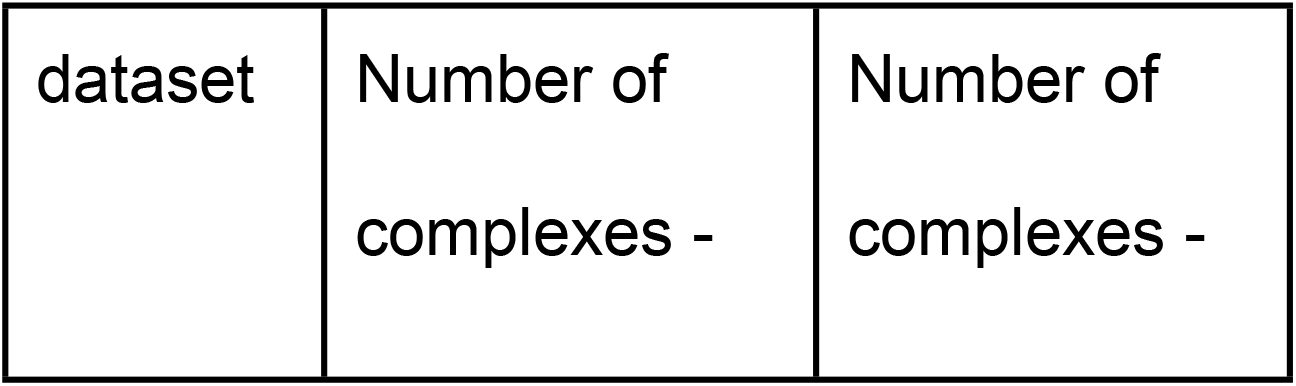
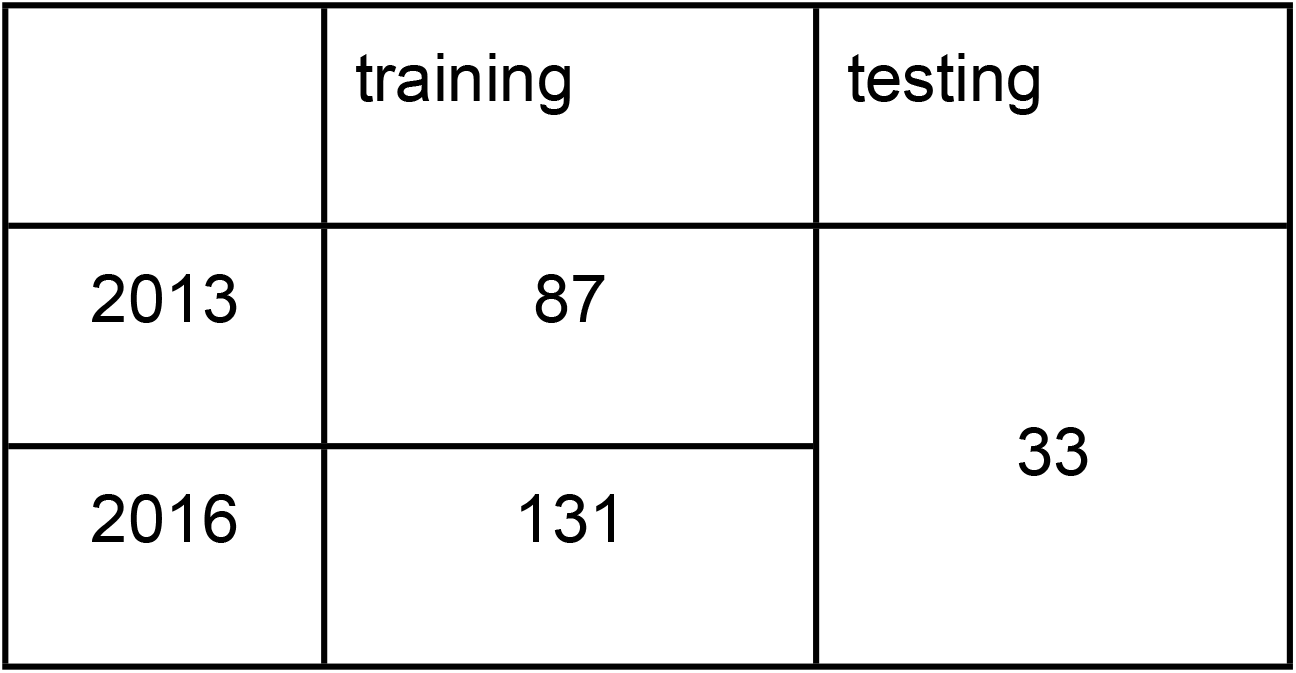
Summary of the 2013 and 2016 datasets.

### Coarse-grained models

To obtain a general model of interactions from the current set of RNA-small molecule complex structures available at the time of the method’s development, we used a coarse-grained representation of both interacting partners. For RNA molecules, we employed a coarse-grained representation used successfully in the SimRNA simulation method, with five ‘beads’ per ribonucleotide residue with positions corresponding to real atoms [53]. In the current version of the scoring function, only the canonical A, G, C, U residues were taken into account. Modified residues (e.g., due to post-transcriptional modification) were not used for developing the potential and were only considered as a steric hindrance. Hence, for docking of ligands to RNA molecules that may have modified residues in the binding sites, we recommend to model the replacement of modified residues with their canonical counterparts, e.g., using the ModeRNA method ([54], also available as a web server, http://iimcb.genesilico.pl/modernaserver/, [55]).

For small-molecule ligands, we applied the concept of pharmacophores, according to the implementation proposed by Taminau et al. [52]. We used six types of pharmacophores and Euclidean vectors derived for each pseudoatom indicating the direction of the given pharmacophore (See: Fig 8). The centers of the HDON, HACC, POSC, and NEGC points coincide with the position of the heavy atom that is labeled as a hydrogen bond donor, acceptor, carrying a positive or a negative charge. The AROM point is positioned in the center of the aromatic ring it represents. The LIPO point is generated in a multistep procedure, where adjacent lipophilic regions are averaged to a single pharmacophore with respect to their lipophilic contribution. If more than one possible pharmacophore can be assigned to a single chemical group, e.g., hydrogen bond donor and acceptor for an amine group, both pharmacophore features are assigned independently.

**Fig 8.**
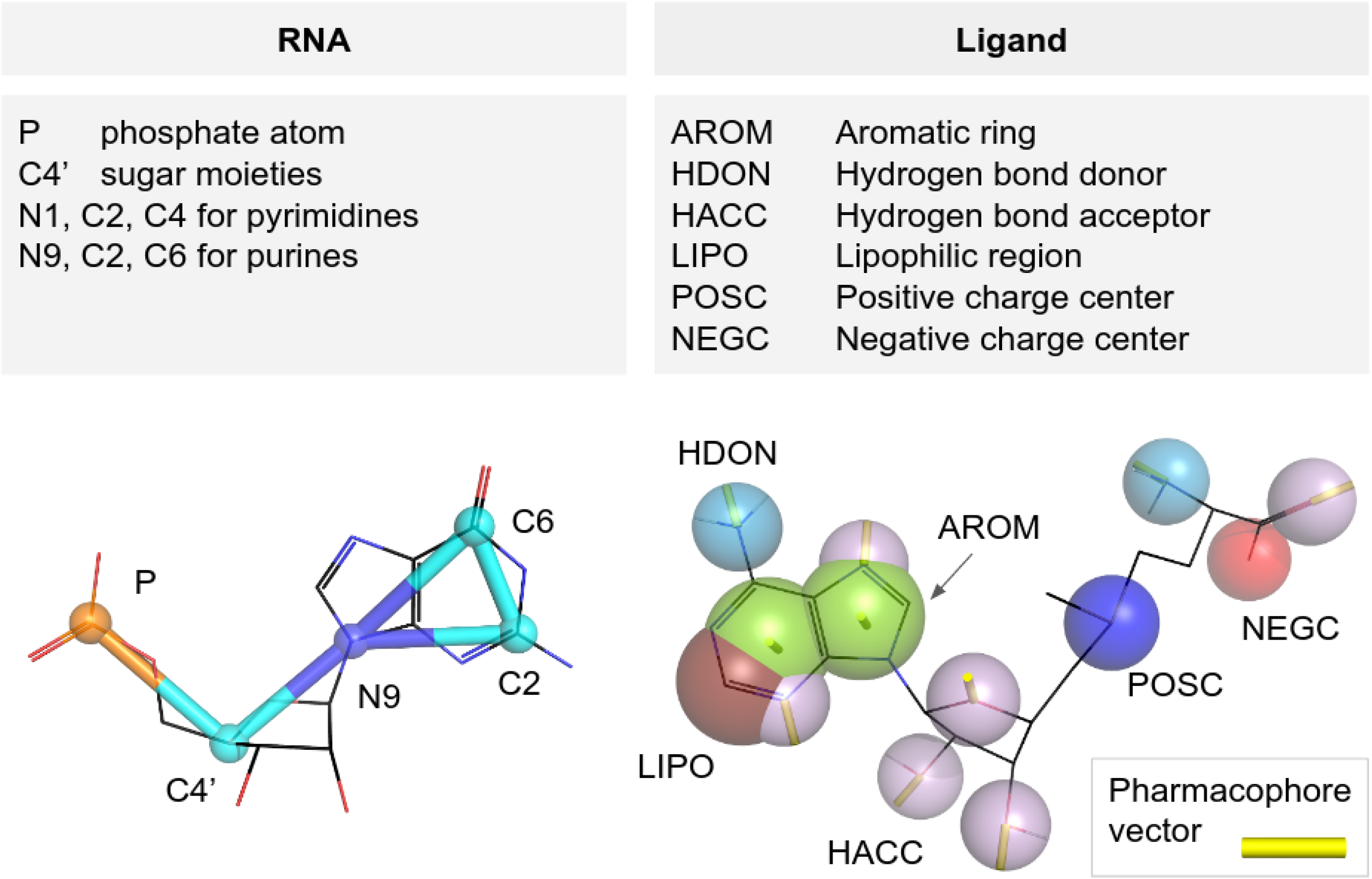
Atoms and pseudoatoms used in the coarse-grained representation of RNA (upper left pane) and ligand molecules (upper right pane), an example of a ribonucleotide (guanosine monophosphate) in SimRNA representation (bottom left) and a small molecule - S-adenosylmethionine (bottom right) in the pharmacophore representation.

### Descriptors of RNA-Ligand interactions

The set of descriptors collected for each RNA-ligand pseudoatom pair consisted of five features that describe geometric relationships between the RNA component and the small molecule component: the distance, the angle between the two selected pseudoatoms of the RNA and the pharmacophore, the angle between the two selected pseudoatoms of the RNA and the pharmacophore vector, the angle between the nucleotide base plane and the pharmacophore, and the angle between the base plane and the pharmacophore vector. The nucleotide plane is defined by atoms P, C4', N9 and C2, C6, N9 for purines and P, C4, N1 and C2, C4, N1 for pyrimidines. The interaction threshold was set at 10 Å, a value identified during the optimization of the scoring function (see: S1 Text). Descriptors collected for each contact were assigned a class depending on the origin of the input ligand’s structure (“positive” or “negative”).

### Machine learning models of interactions

To derive the scoring models from the collected multidimensional interactions statistics, we used supervised machine learning methods. We tested five widely used algorithms (Deep Learning - multi-layer feedforward artificial neural network - DL, Gaussian Naïve Bayes - GNB, k Nearest Neighbors - *k*NN, Random Forests - RF, and Support Vector Machines with RBF kernel - SVM). After benchmarking these methods, (see: S1 Text, chapter “Optimization of parameters of the scoring function”) we selected two best-performing methods: *k*NN - *k* nearest neighbors method and the Deep Learning (multi-layer feedforward artificial neural network).

#### Model implementation details

Machine learning models for each contact group (base - RNA atom - ligand pseudoatom) were derived independently. To remove contact groups with a statistically insufficient number of contacts, we removed 20 out of 120 possible groups with the number of collected interactions below the arbitrary set threshold of 10 “positive” contacts. For the list of contact groups for which machine learning models were built and used for scoring, see Table S2 in S1 Text).

#### Model development and testing

During the development of machine learning models, we used only the training datasets. We applied a cross-validation scheme with five splits of data according to the ligand structural class. Firstly, ligands were clustered into five groups, using RDKit fingerprint Tanimoto similarity and *k*-Medoids clustering method, representing five chemical classes of compounds: 1) amino acids and carboxylic acids, 2) heterocycles and polycyclic compounds, 3) polysugars, 4) amines, 5) alcohols and polyols. Each variant of AnnapuRNA scoring function was trained using data from four clusters and tested on the data coming from the remaining cluster. This training and testing procedure was repeated for all five combinations of clusters, and the scoring performance was averaged. At each cross-validation, split data were class-balanced, i.e., both classes were represented by an equal number of cases.

All descriptors were rescaled to the [0; 1] range. To remove class label noise, for each contact group, we used the Edited Nearest Neighbors algorithm (ENN, [56]) implemented in the imbalanced-learn python package [57]. Parameters of machine learning models were optimized individually for each contact group using Grid Search method. We used two python implementations of machine learning algorithms: scikit-learn (version 0.17, [58]) and H2O (version 3.9.1.3501, https://www.h2o.ai/).

For the optimized set of parameters and developed processing pipeline, we built the final four versions of predictive models - for two selected machine learning algorithms (*k*NN and Deep Learning) using two training datasets (“2013” and “2016”).

### Potential definition

The total score for RNA-Ligand complex is a sum of two terms:

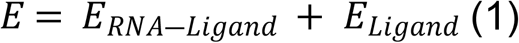

where *E* is the final score of the complex, *E*_*RNA-Ligand*_ is the score of RNA-Ligand interaction and *E*_*Ligand*_ is a score of the internal ligand’s conformation. The score of RNA-Ligand interaction is expressed as a sum of probabilities *p* of a given interaction for all interactions between RNA atoms and ligand pseudoatoms within the cutoff distance 10 Å (Eq. 2 and Fig 9). The probability of RNA-ligand interactions *p* is calculated from a Machine Learning model and it expresses the likelihood that the respective interaction belongs to the “positive” class

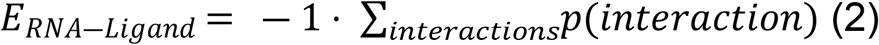

**Fig 9.**
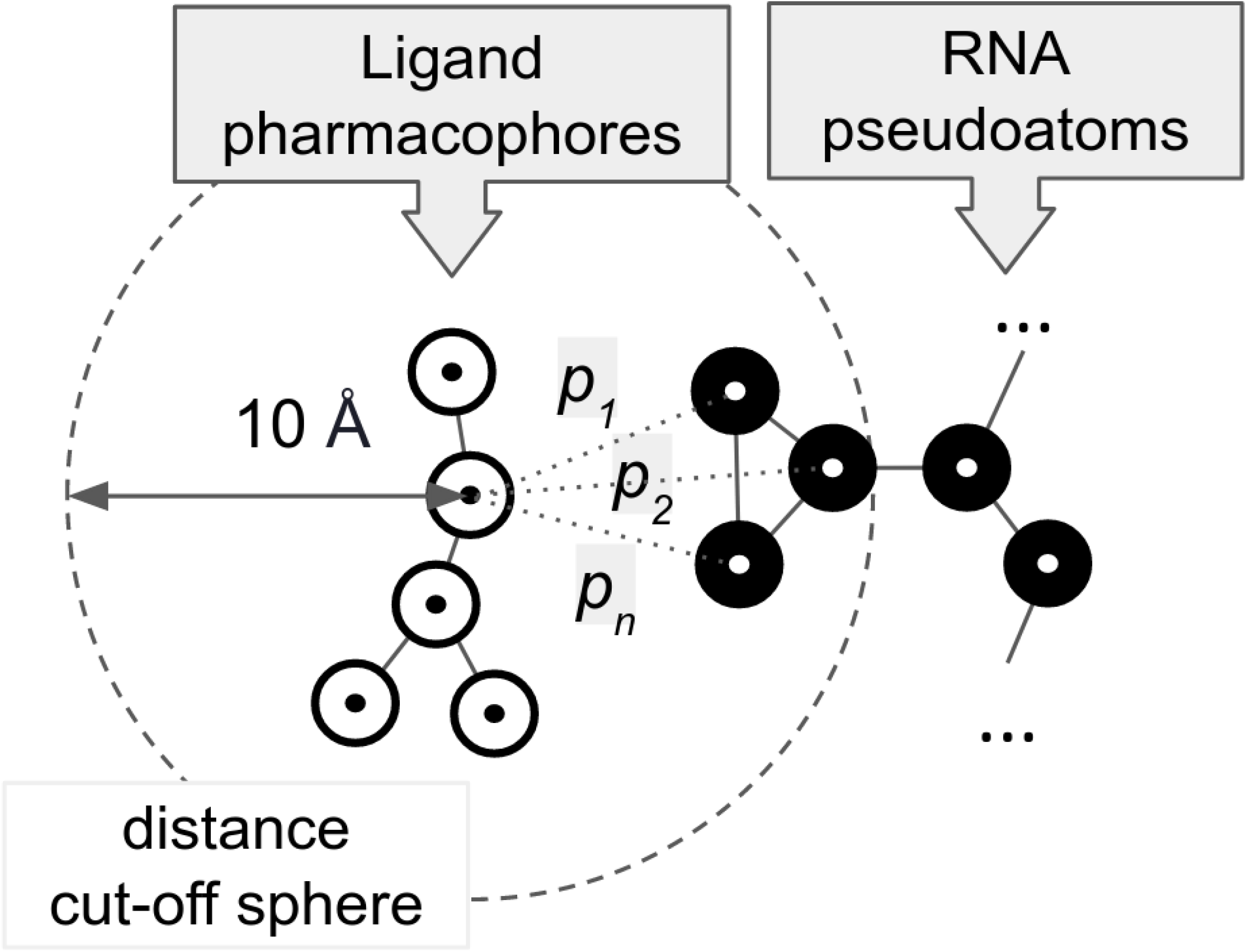
AnnapuRNA scoring function calculates the probabilities *p*_*1*_...*p*_*n*_ of interactions between ligand pharmacophores (white circles) with RNA pseudoatoms (black circles) within the distance of 10 Å. The total score is calculated as a negative sum of all probability values *p* calculated for the given ligand.

The score of internal energy of ligand, *E*_*Ligand*_, is derived from GAFF internal energy of the ligand [59] and is calculated from:

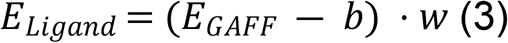

The ligand internal energy contribution is scaled by the factor *b*, which shifts the positive values of the GAFF energy to the negative values. The value of this parameter was set to 473.58, which is equivalent to the third quartile of GAFF energy values calculated for the diversified set of experimentally determined ligand structures deposited in the PDB database. The ligand’s contribution to the final complex score is scaled by the weighting factor *w*. This parameter was set to 0.1 after optimization in a cross-validation experiment but may be changed by the user via a command-line switch.

For the full optimization procedure of parameters of the AnnapuRNA scoring function, see chapter “Optimization of parameters of the scoring function” in S1 Text.

### Construction of the test set

To compare the AnnapuRNA scoring function with the previously published methods, we performed a redocking experiment. The testing set consisted of 33 RNA-small molecule complex structures, included in the set used earlier by Philips et al. [35] and for which we were able to obtain docking results using at least two methods.

Many factors may have an influence on docking performance. Among the most obvious ones are the docking program and the scoring function used. Within this work, we tested three docking programs: Autodock Vina [23], iDock (version 2.2.1, [25]) and rDock with two desolvation potentials: dock and dock_solv (version 2013.1, [30]. For each of the docking programs, we specified a docking volume of 10 Å around the ligand (i.e., the grid box size for AutoDock Vina and iDock was set to 20 Å with a center in the center of mass of the ligand; for rDock, the radius parameter was set to 10 Å). For each ligand, 100 poses were generated (except AutoDock Vina, for which the maximum number of poses is 20).

Generated poses were rescored using nine scoring functions: LigandRNA (in two versions: the one which is described in the original publication, called “2013”, and the second, with an updated potential, called “updated”), rDock (with two potentials: dock and dock_solv), RF-Score-VS (version v2, [60]) and four variants of the scoring function, AnnapuRNA, presented in this work. The four variants correspond to two versions of the scoring function: *k*NN and DeepLearning, each trained on two datasets: “2013” and “2016”. In the final comparison for iDock and AutoDock Vina, we also included results from the scoring function used by the docking program (termed “Internal SF”).

In the real-life scenario of docking small molecules to a macromolecular receptor, an experimentally solved structure of the macromolecule and a *de novo* generated three-dimensional structure of a ligand are used. In our redocking experiment, we wanted to simulate this process and use not only the native ligand structure extracted from the solved complex but also ligand structures generated from one-dimensional SMILES structures. For this purpose we used two widely used tools: OpenBabel with --gen3D option (version 2.3, [49]) and Balloon (version 1.6.2.1236, [61]).

### Measuring the docking performance

To test the performance of our method as well as other methods in their ability to identify the most accurate binding poses, we used the *S*(X) metric [35], which reports the lowest RMSD to the reference ligand among the top X scoring poses. Here we used three variants of this metric: *S*(1), *S*(3), and *S*(5), which report the RMSD of the top-scoring pose, the top three poses, and the top five poses, respectively. The RMSD between the docked ligand pose and the reference (experimentally determined) ligand pose was calculated using an in-house python script, and it took into account the symmetry within the ligand structure. We employed the Wilcoxon signed-rank to test if the median scores obtained from the AnnapuRNA algorithm was significantly different (in our case: better) than for other scoring functions and reported two-tailed *p* values. Statistical analysis was performed in the KNIME Analytics Platform 4.1.3 with a Wilcoxon Signed-Rank node [62]. Correlation coefficients were calculated with pandas 1.0.5 [63].

## Acknowledgments

We thank Anna Philips for help with LigandRNA and other members of the Bujnicki laboratory for their help. We thank Katarzyna Merdas and J. Mahita for critical reading and proof-reading of the manuscript. We also thank the StackOverflow community (https://stackoverflow.com/) for support and building a useful knowledge resource.

---

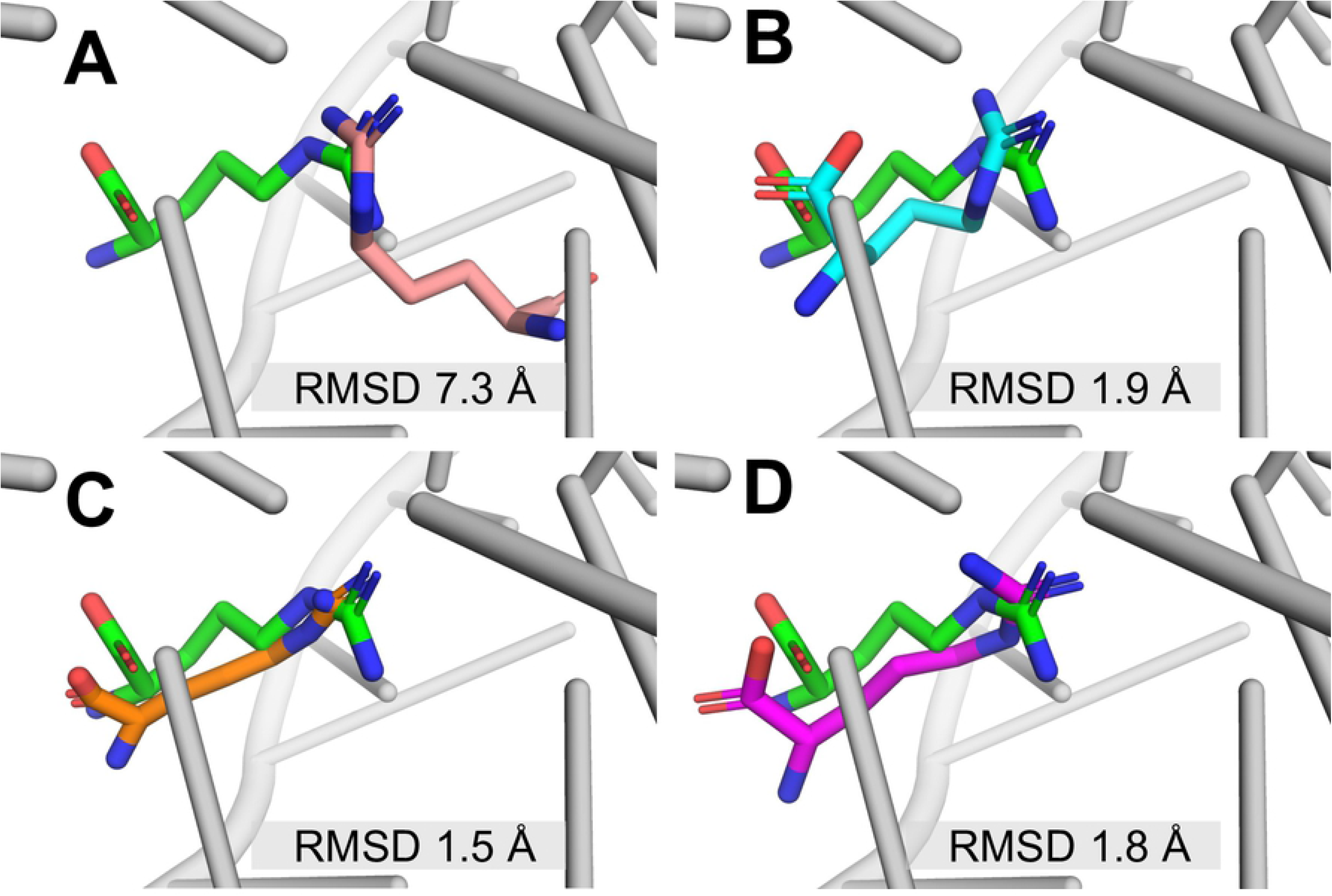

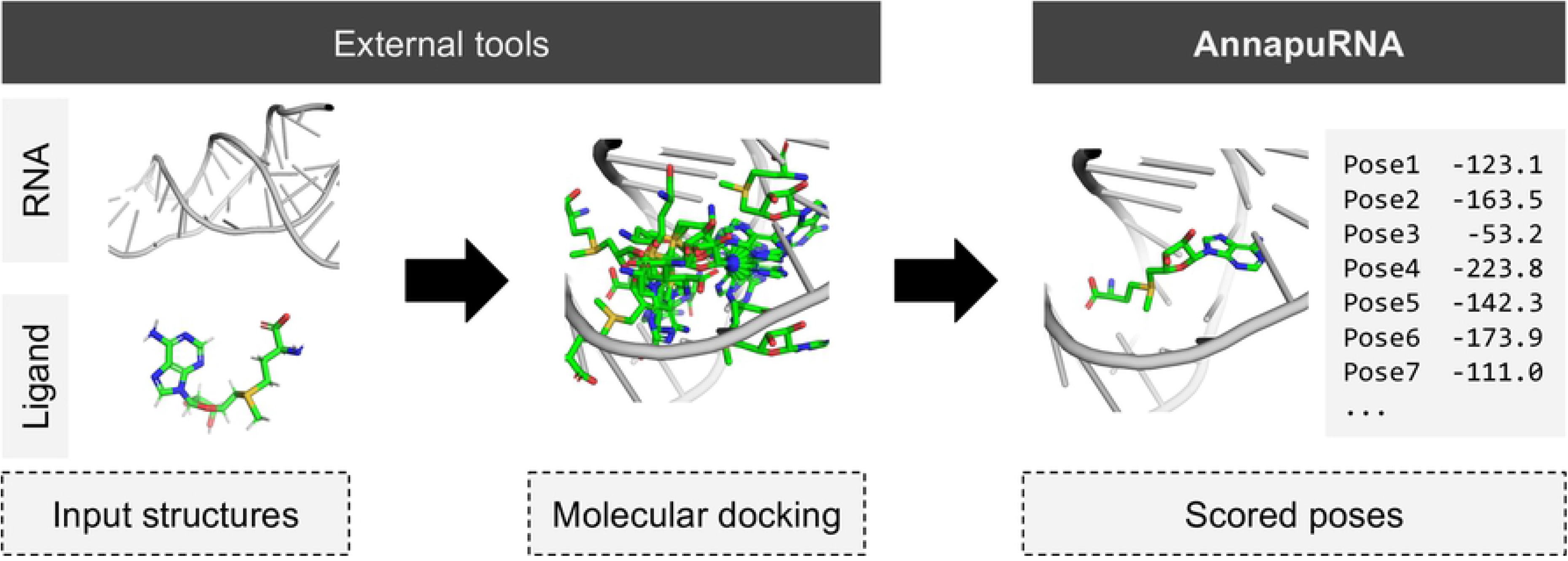

